# Rapid identification of homozygosity and site of wild relative introgressions in wheat through chromosome-specific KASP genotyping assays

**DOI:** 10.1101/633842

**Authors:** Surbhi Grewal, Stella Hubbart-Edwards, Caiyun Yang, Urmila Devi, Lauren Baker, Jack Heath, Stephen Ashling, Duncan Scholefield, Caroline Howells, Jermaine Yarde, Peter Isaac, Ian P. King, Julie King

## Abstract

For future food security it is important that wheat, one of the most widely consumed crops in the world, can survive the threat of abiotic and biotic stresses. New genetic variation is currently being introduced into wheat through introgressions from its wild relatives. For trait discovery, it is necessary that each introgression is homozygous and hence stable. Breeding programs rely on efficient genotyping platforms for marker-assisted selection (MAS). Recently, single nucleotide polymorphism (SNP) based markers have been made available on high-throughput Axiom^®^ SNP genotyping arrays. However, these arrays are inflexible in their design and sample numbers, making their use unsuitable for long-term MAS. SNPs can potentially be converted into Kompetitive allele-specific PCR (KASP™) assays which are comparatively cost-effective and efficient for low-density genotyping of introgression lines. However, due to the polyploid nature of wheat, KASP assays for homoeologous SNPs can have difficulty in distinguishing between heterozygous and homozygous hybrid lines in a backcross population. To identify co-dominant SNPs, that can differentiate between heterozygotes and homozygotes, we PCR-amplified and sequenced genomic DNA from potential single-copy regions of the wheat genome and compared them to orthologous copies from different wild relatives. A panel of 620 chromosome-specific KASP assays have been developed that allow rapid detection of wild relative segments and provide information on their homozygosity and site of introgression in the wheat genome. A set of 90 chromosome-nonspecific assays was also produced that can be used for genotyping introgression lines. These multipurpose KASP assays represent a powerful tool for wheat breeders worldwide.

## Introduction

Bread wheat (*Triticum aestivum* L.) is one of the most widely grown crops in the world and accounts for almost one-fifth of the human calorie intake (FAO, 2017). Its allohexaploid (AABBDD; 2n = 6× = 42) genome was derived from the hybridisation of diploid *Aegilops tauschii* (DD; 2n=2×=14) with tetraploid *Triticum turgidum* ssp. *dicoccoides* (AABB; 2n=4×=28) (Dubcovsky and Dvorak, 2007; Matsuoka, 2011). Due to this hybridisation event, followed by domestication and inbreeding, genetic variation has reduced in modern cultivated wheat (Haudry et al., 2007). However, genetic diversity is crucial if the wheat species is to survive and adapt to the threat of abiotic and biotic stresses. It has been suggested that interspecific crossing of wheat with its wild relatives can enrich wheat’s gene pool with novel diversity (Reynolds et al., 2011). One strategy, recently called ‘introgressiomics’ (Prohens et al., 2017), consists of a whole-genome introgression approach involving transfer of chromosome segments from the entire genome of a wild relative species into the wheat background, irrespective of any traits that the wild relative might carry and a number of such studies have already been undertaken (Grewal et al., 2018a; Grewal et al., 2018b; King et al., 2018; King et al., 2017; Valkoun, 2001). In this pre-breeding strategy, the interspecific hybrids are repeatedly backcrossed to the elite wheat parent to reduce the number and size of the introgressed segments and self-fertilised to obtain stable homozygous introgressions that can be utilised for trait analysis (King et al., 2019). Previously, wild relative introgressions were detected using labour-intensive cytogenetic techniques (Lukaszewski et al., 2005). More recently, molecular markers provide high throughput and cost-effective evaluation of introgressions in large numbers of lines (Thomson, 2014).

Some studies have used co-dominant markers such as simple sequence repeats (SSRs) to detect wild relative introgressions in wheat (Adonina et al., 2004; Qi et al., 2010; Quarrie et al., 2005; Rodríguez-Suárez et al., 2011; Zhao et al., 2013). However, with these being cost-ineffective, laborious and time-consuming to use, they have limited potential in wheat breeding programmes. Single nucleotide polymorphism (SNP) markers, on the other hand, have now become common place in wheat genotyping (Akhunov et al., 2009; Bevan and Uauy, 2013; Davey et al., 2011) and marker assisted selection (MAS). However, in polyploid species such as wheat, the development of SNP markers has been challenging due to the presence of homoeologous and paralogous copies of genes (Edwards et al., 2009; Kaur et al., 2012) and distinguishing between interspecific SNPs from intergenomic polymorphisms within wheat can be complicated and error-prone (Akhunov et al., 2009). Exome-based sequencing has provided a huge resource of SNPs between wheat varieties (Allen et al., 2013; Winfield et al., 2012). Many of these have been developed into high-density SNP arrays (Allen et al., 2017; Rimbert et al., 2018; Wang et al., 2014; Winfield et al., 2016) for high-throughput genotyping in wheat. An Axiom^®^ Wheat Relative SNP Genotyping Array has also been developed and used in studies for identification and characterisation of wild relative introgressions in a wheat background (Grewal et al., 2018a; Grewal et al., 2018b; King et al., 2018; King et al., 2017). Although these powerful SNP genotyping platforms can be ultra-high-throughput and efficient, their use in crop breeding has been limited because they are inflexible in their design and use (Rasheed et al., 2017). This leaves wheat breeders who want to carry out medium-to low-density genotyping on large number of plants with very few options.

More recently, the Kompetitive allele-specific PCR (KASP™) system has been demonstrated to be a more flexible, efficient and cost-effective system for genotyping in wheat (Allen et al., 2013; Neelam et al., 2013; Tan et al., 2017; Yu et al., 2017) and other crop species (Semagn et al., 2014; Steele et al., 2018; Zhao et al., 2017). The KASP system allows 1) conversion of SNPs from fixed chip platforms to a stand-alone format where hundreds to thousands of samples can be genotyped with relatively fewer markers and 2) flexibility of customisation of the genotyping run with different combinations of SNP markers and sample numbers. But this technology has two major drawbacks for wheat-wild relative genotyping. Firstly, it requires the identification and characterisation of interspecific SNPs among an excess of homoeologous and paralogous SNPs. Secondly, as this platform was primarily developed for diploid species, there are problems with the scoring of interspecific SNPs in polyploid heterozygotes, such as segregating backcross populations. For the KASP system to detect a wild relative segment in an allohexaploid wheat background it has to accurately distinguish between different call ratios (Allen et al., 2011; Allen et al., 2013). For example, if the KASP assay is for a SNP which has three homoeologous copies in wheat, it will be extremely difficult to distinguish between a heterozygous introgression having a call ratio of 5:1 and a homozygous introgression having a call ratio of 4:2, in a self-fertilised backcross line (Allen et al., 2011). In contrast, if the SNP assay is for a SNP which amplifies only a single homoeologous/paralogous copy in wheat (co-dominant), then this system would be easily capable of differentiating between a heterozygous (call ratio of 1:1) and a homozygous (call ratio of 2:0) introgression in a segregating population.

A recent study successfully converted a panel of PCR markers to KASP™ markers for functional genes in wheat (Rasheed et al., 2016). A number of array-based, putative co-dominant SNPs have been reported for various wild relatives (Grewal et al., 2018a; Grewal et al., 2018b; King et al., 2018; King et al., 2017) which could potentially be converted into KASP™ assays. However, it is difficult to design polymerase chain reaction (PCR) primers for array-based probes due to the high level of sequence polymorphism between wheat and its wild relatives. It is also possible that homoeologous sequences that may not have bound to array-based probes due to sequence divergence could be amplified by the KASP primers. However, careful primer design can lead to the successful amplification of just one homoeolog/paralog. Moreover, targeting single-copy regions of the wheat genome for SNPs could be a more fruitful strategy since these sequences, by definition, will not have homoeologous copies and thus, should not suffer from the interference usually encountered.

In this study, we have exploited chromosome-specific sequences in wheat, i.e., sequences that are found only on a particular chromosome of wheat, for SNPs with wild relative species. Some of these SNPs were subsequently converted to KASP assays and where a target SNP sequence was not chromosome-specific, i.e. having other homoeologous copies, the KASP assays were designed to potentially amplify only the target subgenome of wheat. This work has resulted in 620 chromosome-specific co-dominant KASP assays, evenly spread across the hexaploid wheat genome. These assays will allow rapid identification of homozygosity of wild relative introgressions and their site of recombination within wheat, in segregating populations. In addition, a set of 90 chromosome-nonspecific KASP assays is also reported that are useful for genotyping lines for the presence of wild relative segments. Validation was carried out through genotyping backcross populations of these wild relative species and genomic *in situ* hybridisation (GISH). These KASP assays are valuable tools in wheat-wild relative introgression studies and are predicted to be useful for detection of many other wheat wild relatives which will be of considerable interest to the wheat research community.

## Results

### SNP Discovery

A BLASTN search of all the 36,711 SNP-containing probe sequences on the Axiom^®^ Wheat-Relative Genotyping Array against the wheat reference sequence, IWGSC CSS v3 (IWGSC, 2014), resulted in 2,716 probes that had a BLAST hit to only 1 contig (**Table 1**). From these 2,716 target SNP-containing probe sequences, it was possible to design primers for 2,170 from the flanking 500 bp genomic sequence. Genomic DNA from two wheat varieties, Paragon and Chinese Spring, along with ten wild relatives of wheat, *Amblyopyrum muticum, Thinopyrum bessarabicum, Thinoypyrum intermedium, Thinopyrum elongatum, Thinopyrum ponticum, Aegilops speltoides, Aegilops caudata, Triticum urartu, Triticum timopheevii* and *Secale cereale*, were used to test the primers for PCR amplification. 1,721 primer pairs were successful in amplifying at least one wheat variety and one wild relative species (**Table 1**). The PCR amplification resulted in 13,731 PCR products that were sent for sequencing. Of these, 61.67% of samples were successfully sequenced. **Table 1** shows the distribution of the number of primer pairs designed and samples sequenced across the 21 chromosomes of wheat.

**Table 1.** Distribution, across wheat chromosomes, of the number of probes on the Axiom^®^ Wheat-Relative Genotyping Array with BLASTN hit to a single contig in the wheat genome, the primers designed from these probes, PCR products sent for sequencing, the SNPs discovered and selected for KASP assay design.

| Wheat<br>Chromosome | Probes on WRGA†<br>chip with blast to<br>1 contig‡ | Primer<br>pairs<br>designed | Primers<br>worked | PCR<br>products<br>sent for<br>sequencing | Successfully<br>sequenced<br>samples | Successfully<br>sequenced<br>samples (%) | SNPs<br>discovered<br>(across all<br>10 species) | SNPs sent<br>for KASP<br>assay design |
| --- | --- | --- | --- | --- | --- | --- | --- | --- |
| 1A | 83 | 63 | 54 | 416 | 276 | 66.35 | 93 | 45 |
| 1B | 141 | 99 | 78 | 507 | 312 | 61.54 | 135 | 56 |
| 1D | 144 | 99 | 83 | 608 | 334 | 54.93 | 154 | 46 |
| 2A | 112 | 91 | 79 | 540 | 396 | 73.33 | 119 | 48 |
| 2B | 159 | 132 | 99 | 605 | 442 | 73.06 | 141 | 54 |
| 2D | 130 | 111 | 93 | 633 | 447 | 70.62 | 126 | 43 |
| 3A | 85 | 86 | 86 | 747 | 278 | 37.22 | 117 | 45 |
| 3B | 294 | 266 | 192 | 1469 | 1009 | 68.69 | 168 | 53 |
| 3D | 79 | 76 | 62 | 469 | 296 | 63.11 | 126 | 45 |
| 4A | 95 | 77 | 75 | 833 | 447 | 53.66 | 96 | 46 |
| 4B | 88 | 73 | 57 | 415 | 224 | 53.98 | 65 | 35 |
| 4D | 73 | 64 | 67 | 586 | 329 | 56.14 | 90 | 43 |
| 5A | 68 | 55 | 46 | 382 | 262 | 68.59 | 78 | 49 |
| 5B | 167 | 122 | 112 | 833 | 530 | 63.63 | 115 | 56 |
| 5D | 155 | 110 | 97 | 729 | 426 | 58.44 | 116 | 47 |
| 6A | 102 | 97 | 83 | 642 | 501 | 78.04 | 113 | 45 |
| 6B | 92 | 84 | 72 | 531 | 348 | 65.54 | 107 | 53 |
| 6D | 140 | 128 | 119 | 897 | 604 | 67.34 | 125 | 49 |
| 7A | 90 | 60 | 49 | 403 | 220 | 54.59 | 73 | 44 |
| 7B | 140 | 89 | 74 | 521 | 301 | 57.77 | 111 | 47 |
| 7D | 279 | 188 | 44 | 965 | 469 | 48.60 | 106 | 51 |
| **Total** | **2716** | **2170** | **1721** | **13731** | **8451** | **61.67** | **2374** | **1000** |

† Axiom® Wild Relative Genotyping Array
‡ BLASTN against IWGSC CSS v3

Orthologous sequences from the wild relative species and at least one wheat variety were aligned to identify putative interspecific SNPs. A SNP was given preference if it was common between multiple wild relatives. A maximum of one SNP/wild relative species from a primer pair was selected to maximise genome coverage. In total, 2,374 putative SNPs, from 8,451 sequences, were obtained across all ten wild relative species (**Table 1**) and their distribution across the wild relative species and the wheat chromosomes is detailed in **Table S1**. The highest number of SNPs were obtained for *Am. muticum* and *Th. bessarabicum* with 458 each while the least number of SNPs were 248 and 254, obtained for *Th. ponticum* and *T. urartu*, respectively.

### Primer design for chromosome-specific assays

The sequence flanking the target SNP was used in a second BLASTN search against the most recent wheat genome sequence, IWGSC Refseq v1 (IWGSC et al., 2018). This additional BLAST search was added to confirm that there were no other homoeologous copies of the target SNP sequences (the improved high-quality reference assembly for wheat had become available after SNP discovery had been completed). This BLASTN search revealed that only 433 of the 2374 (18.2%) SNP-containing sequences had a single-copy in the wheat genome. The rest of the sequences had at least one homoeologue, with 67% of the sequences having a homoeologous copy on all 3 subgenomes of wheat (**Table S2**). The results also showed that 57.5% of the sequences with more than one copy in wheat had homoeologous SNPs i.e. the target SNP was polymorphic between the homoeologues in wheat.

Ideally, once a SNP was identified as having flanking sequence suitable for primer annealing, an allele-specific KASP assay could be designed (**Figure 1A**). However, in cases where there were homoeologous copies of target SNP sequences, primer optimisation was required so that the KASP™ assays would be specific to a particular chromosome in wheat. To do this, a unique base(s) in the flanking sequence of the target subgenome was identified, i.e. a base(s) that was specific to one homoeologue, but also be present in the orthologous wild relative sequence. This single unique base was incorporated into the common primer during the KASP™ assay design to obtain target amplification specificity, also known as primer ‘anchoring’, as shown in **Figure 1B**. If such unique bases could be identified in the target SNP’s flanking sequence, the SNP was categorised as potentially chromosome-specific. If no specific allele was identified in the target subgenome, the SNP was characterised as chromosome-nonspecific. Of the 1,941 sequences that had more than one copy in wheat, 1,488 (76.7%) putative SNPs had the potential for chromosome-specific assays to be designed while 453 (23.3%) putative SNPs could only be selected as chromosome-nonspecific assays (**Table S2**). Where the target SNP was homoeologous, i.e. polymorphic within wheat, it was selected only if the assay designed for it was potentially chromosome-specific (**Figure 1B**).

**Figure 1.**
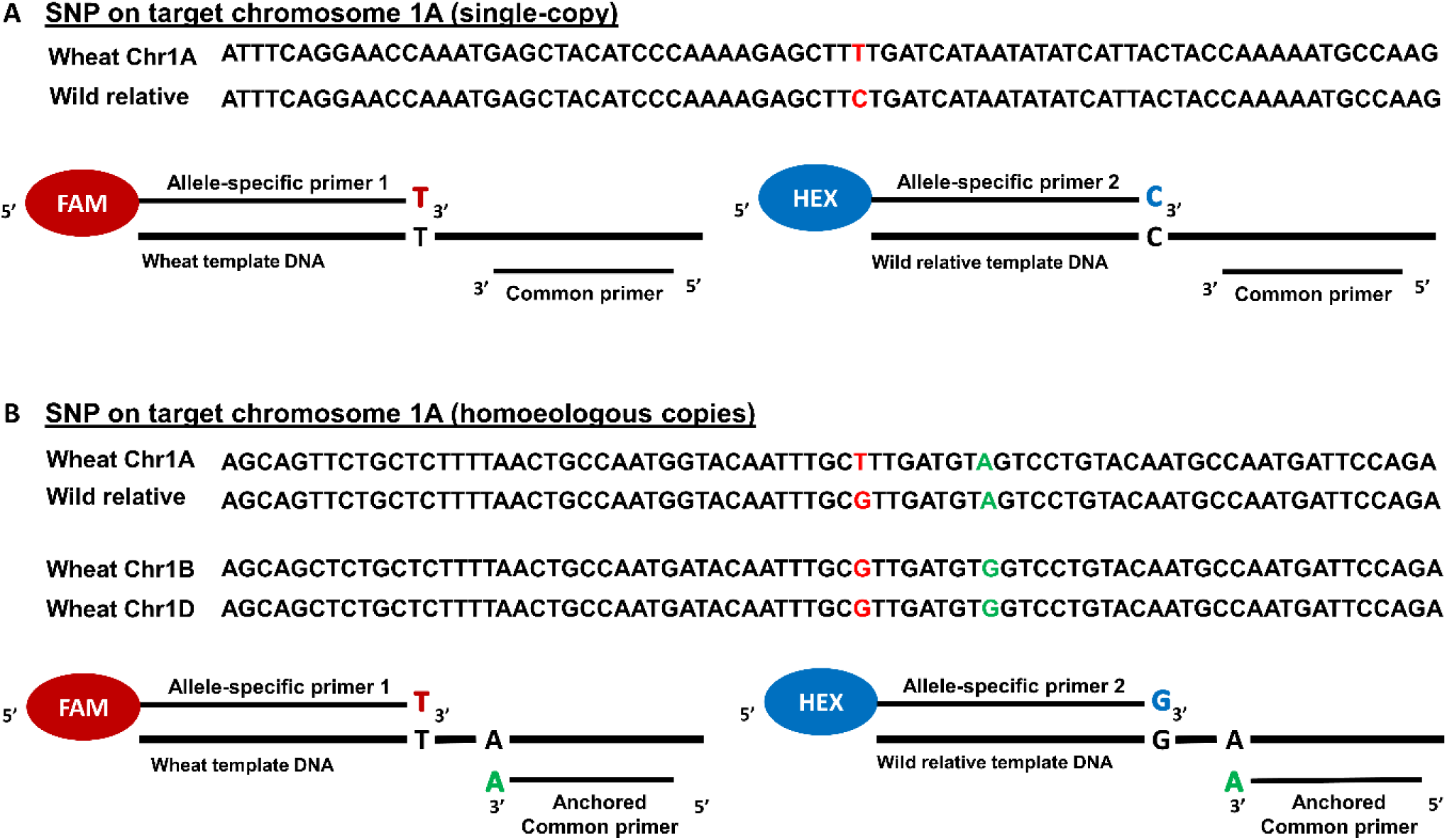
Components of a KASP assay when the target SNP is in a single-copy region of the wheat genome versus when there are homoeologous copies. A) For a target single-copy SNP T/C, on wheat chromosome 1A, the KASP assay mix contains two allele-specific forward primers and one common reverse primer. B) For a target SNP T/G, on wheat chromosome 1A having homoeologous copies on chromosomes 1B and 1D, the KASP assay mix contains two allele-specific forward primers and one common reverse primer with its 3’ end anchored to a base which is unique to the target subgenome in wheat and is absent in the homoeologous copies. The anchored base is also present in the wild relative sequence. The allele-specific primers each harbour a unique tail sequence that corresponds with a universal FRET (fluorescence resonant energy transfer) cassette; one labelled with FAM™ dye and the other with HEX™ dye.

### SNP validation and characterisation

A subset of 1,000 putative SNPs was selected for validation using the KASP™ genotyping platform (**Table 1 and Table S3**) of which 864 were potentially chromosome-specific and selected to be evenly distributed across the wheat chromosomes, wherever possible. To fill in the gaps, 136 potentially chromosome-nonspecific SNPs were selected to make up the total to 1,000 SNPs. The target SNP sequences, along with any annotations for chromosome specificity, were sent to LGC Genomics for KASP™ assay design. The SNP validation, also performed by LGC Genomics, was done through genotyping the parental wheat varieties, wild relative accessions (**Table S4**) and all the Chinese Spring nullisomic lines as controls along with screening 4,666 segregating lines (BCnFn) from the backcrossed and self-fertilised populations of the ten wild relatives (**Table S5**).

Of the 1,000 putative interspecific SNPs, 710 were polymorphic (between all the wheat varieties and at least one wild relative species), 17 were polymorphic within wheat itself (polymorphism between the homoeologous copies), 3 were polymorphic between the four wheat varieties and 270 failed to generate a useful amplification signal. Primers were not redesigned when amplification failed. It was noted that of the failed assays, 141 failed to amplify the target wild relative accessions, i.e. they either only worked for the wheat varieties or were monomorphic with non-target wild relatives. The genotypes obtained for all the parental and nullisomic tetrasomic lines are provided in **Data S1**.

**Figures 2A-J** depict how genotyping with a chromosome-specific and nonspecific KASP assay worked. In **Figure 2A**, the target SNP on chromosome 1A is potentially chromosome-specific (either because the SNP was only present on a single locus in wheat or the primer design was optimised with primer anchoring as shown in **Figure 1B**) where the wheat allele is T/T and the wild relative allele is C/C. Screening a line having no wild relative introgression with this assay resulted in a wheat call (T/T; **Figure 2B**). Screening introgression lines with this assay resulted in three separate clusters for homozygote and heterozygote individuals. When a line had a heterozygous wild relative introgression, this KASP assay gave a heterozygous call (C/T; **Figure 2C**) but if a line had a homozygous wild relative introgression, the assay resulted in the wild relative call (C/C; **Figure 2D**). The chromosome-specificity was validated when the KASP assay was used to genotype the corresponding chromosome’s nullisomic tetrasomic line (N1AT1B) and resulted in a null call as shown in **Figure 2E**. Screening the same population with chromosome-nonspecific SNP assays produced a more scattered cluster where homozygous and heterozygous loci were indistinguishable. For example in **Figure 2F**, a chromosome-nonspecific KASP assay for a non-homoeologous SNP (not polymorphic between the homoeologues in wheat), produced a heterozygous call (C/T; **Figure 2G**) even if the line had a homozygous wild relative introgression and resulted in a wheat call (T/T; **Figure 2H**) in the corresponding nullisomic tetrasomic line N1AT1B (due to the presence of the allele on chromosomes 1B and 1D in both cases). Amongst the 710 validated polymorphic markers, 620 (87%) appeared to be chromosome-specific in wheat capable of distinguishing between homozygous and heterozygous lines (**Table 2**). The remaining 90 KASP™ assays were chromosome-nonspecific i.e. the SNP was present on more than one homoeologue but not polymorphic between them.

**Figure 2.**
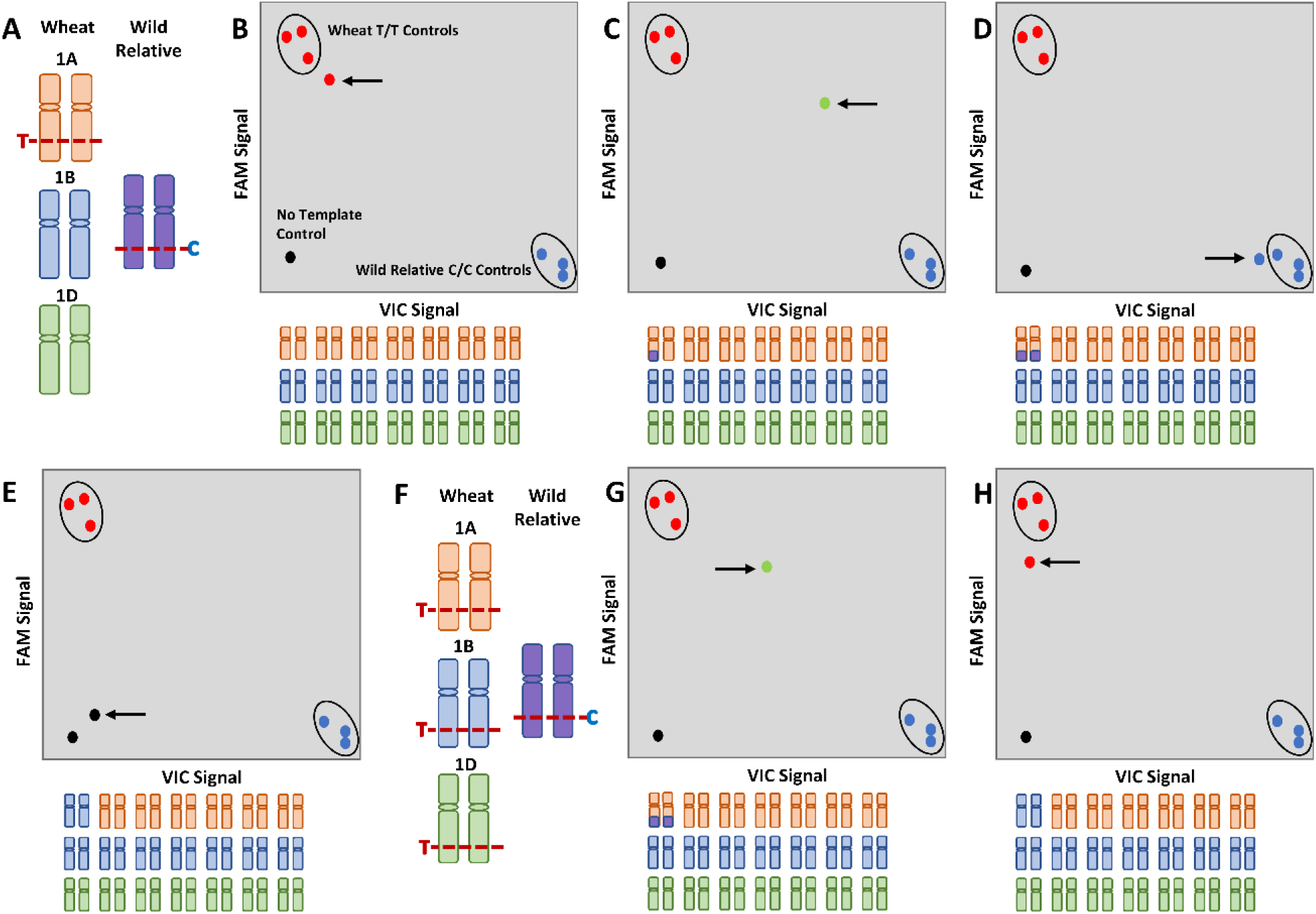
Illustration of genotyping when using chromosome-specific and nonspecific KASP assays. A) A chromosome-specific SNP T/C, on wheat chromosome 1A, used for KASP assay design and genotyping of B) a line with no wild relative introgression shows a homozygous wheat call (red circle indicated by arrow), C) a line with a heterozygous introgression shows a heterozygous call (green circle indicated by an arrow), D) a line with a homozygous introgression shows a homozygous wild relative call (blue circle indicated by an arrow) and E) a nullisomic tetrasomic N1AT1D line shows a no call (black circle indicated by an arrow). F) A chromosome-nonspecific SNP T/C, having homoeologous copies on wheat chromosomes 1A, 1B and 1D, used for KASP assay design and genotyping of G) a line with a homozygous introgression shows a heterozygous call (green circle indicated by an arrow) and H) a nullisomic N1AT1D line shows a homozygous wheat call (red circle indicated by an arrow). In all scenarios, the wheat positive controls are genotyped as T/T (red circles), the wild relative positive controls are genotyped as C/C (blue circles) and the no template control is genotyped as no call (black circle).

**Table 2.** Distribution, across the wheat chromosomes, of the number of chromosome-specific KASP assays validated as diagnostic for each of the ten wild relative species used in this work. Since many of the chromosome-specific assays on a chromosome are diagnostic for more than one wild relative species, the last column indicates the number of unique assays that were validated for each chromosome.

| Wheat<br>Chromosome | *Th.*<br>*bessarabicum* | *Ae.*<br>*caudata* | *Th.*<br>*elongatum* | *Th.*<br>*intermedium* | *Am.*<br>*muticum* | *Th.*<br>*ponticum* | *S.*<br>*cereale* | *Ae.*<br>*speltoides* | *T.*<br>*timopheevii* | *T.*<br>*urartu* | Unique<br>Assays |
| --- | --- | --- | --- | --- | --- | --- | --- | --- | --- | --- | --- |
| 1A | 10 | 10 | 12 | 14 | 5 | 13 | 4 | 5 | 11 | 10 | 29 |
| 1B | 8 | 10 | 17 | 16 | 10 | 20 | 14 | 15 | 15 | 1 | 36 |
| 1D | 9 | 11 | 18 | 19 | 8 | 19 | 4 | 5 | 7 | 2 | 30 |
| 2A | 8 | 8 | 15 | 14 | 11 | 16 | 7 | 2 | 16 | 16 | 33 |
| 2B | 14 | 11 | 21 | 19 | 13 | 24 | 10 | 20 | 25 | 3 | 41 |
| 2D | 10 | 8 | 15 | 19 | 11 | 16 | 3 | 6 | 9 | 3 | 28 |
| 3A | 4 | 3 | 8 | 11 | 3 | 10 | 7 | 3 | 10 | 12 | 27 |
| 3B | 10 | 11 | 19 | 18 | 10 | 17 | 6 | 14 | 18 | 3 | 36 |
| 3D | 6 | 8 | 14 | 12 | 5 | 12 | 7 | 4 | 4 | 2 | 22 |
| 4A | 7 | 8 | 18 | 18 | 5 | 20 | 9 | 8 | 12 | 12 | 31 |
| 4B | 3 | 6 | 9 | 11 | 11 | 9 | 3 | 11 | 14 | 0 | 25 |
| 4D | 4 | 6 | 18 | 18 | 7 | 13 | 5 | 5 | 5 | 2 | 26 |
| 5A | 8 | 10 | 14 | 14 | 2 | 14 | 10 | 4 | 11 | 11 | 28 |
| 5B | 10 | 14 | 15 | 20 | 6 | 16 | 9 | 17 | 18 | 2 | 37 |
| 5D | 11 | 12 | 18 | 16 | 13 | 19 | 4 | 6 | 10 | 5 | 31 |
| 6A | 10 | 4 | 12 | 15 | 6 | 13 | 5 | 4 | 10 | 10 | 28 |
| 6B | 6 | 7 | 18 | 11 | 7 | 14 | 4 | 11 | 16 | 3 | 31 |
| 6D | 8 | 5 | 8 | 10 | 10 | 9 | 6 | 2 | 7 | 4 | 19 |
| 7A | 5 | 4 | 10 | 7 | 6 | 9 | 3 | 1 | 11 | 9 | 21 |
| 7B | 8 | 4 | 15 | 20 | 11 | 18 | 6 | 9 | 10 | 1 | 32 |
| 7D | 7 | 13 | 9 | 19 | 9 | 13 | 5 | 2 | 10 | 3 | 29 |
| **Total** | **166** | **173** | **303** | **322** | **169** | **314** | **131** | **154** | **249** | **114** | **620** |

Some assays were validated for species in which the SNP hadn’t been detected during SNP discovery. For example, *Th. intermedium* was found to have 322 working chromosome-specific assays (**Table 2**) although only 255 SNPs were selected for assay design (**Table S3**). *T. urartu* had the least number of chromosome-specific assays polymorphic with wheat with 114 spread across the wheat genome and none polymorphic with chromosome 4B (**Table 2**). Many of the KASP™ assays were diagnostic for more than one wild relative species. **Table S6** shows the number of chromosome-specific assays common between the different wild relatives. The various Thinopyrum species had more assays common between them than with other wild relative species indicating sequence conservation within the Thinopyrum genus. **Data S2** shows which wild relative species, each of the 710 validated KASP assays are diagnostic for. More than half of the assays (370 assays) were polymorphic between wheat and at least 3 different wild relative species. The physical location of all the polymorphic SNPs, chromosome-specific and nonspecific, and their distribution in the wheat genome is represented in **Figure 3.**

**Figure 3.**
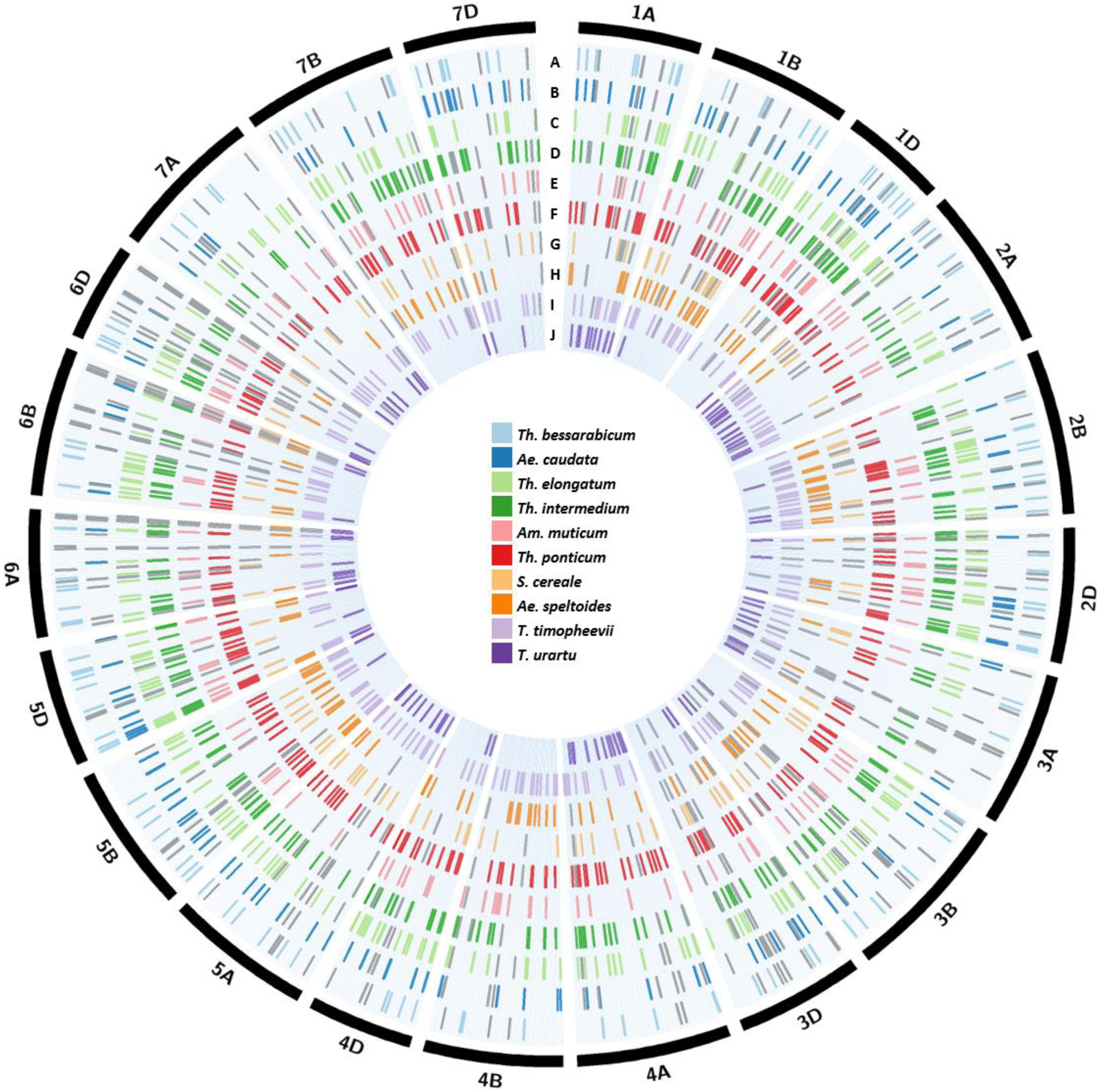
Circular representation of the physical distribution of the chromosome-specific and nonspecific SNPs, for ten wild relative species, across all wheat chromosomes. A coloured line in each ring represents the physical position, on the wheat chromosome (obtained after BLASTN analysis against the IWGSC RefSeqv1 assembly of the wheat genome; IWGSC et al., 2018), of a SNP polymorphic between wheat and A) *Th. bessarabicum*, B) *Ae. caudata*, C) *Th. elongatum*, D) *Th. intermedium*, E) *Am. muticum*, F) *Th. ponticum*, G) *S. cereal*e, H) *Ae. speltoides*, I) *T. timopheevii* and J) *T. urartu*. SNPs for chromosome-specific KASP assays are shown in the colour designated to the wild relative species while SNPs for chromosome-nonspecific assays are shown in grey. The latter are marked for each homoeologous copy in the wheat genome.

BLASTN analysis showed that 368 of the 620 (62%) chromosome-specific assays were from single-copy regions of the wheat genome (**Data S3**) while the rest had at least one other homoeologous copy. However, due to primer anchoring, the latter only amplified the target chromosomes and were thus classified as chromosome-specific. A BLASTX search of the chromosome-specific SNP-containing sequences against the annotated wheat reference sequence Refseq v1 showed that 275 KASP assays were in protein coding regions. Of these, 145 (52%) loci were in single-copy regions and the remaining 130 had more than one homoeologue in wheat (**Data S3)**. The BLASTN results of the chromosome-nonspecific assays are shown in **Data S4**.

### Genotyping with chromosome-specific markers and validation by genomic *in situ* hybridisation (GISH)

In addition to the wheat varieties and the wild relative accessions, the KASP markers were also used to genotype a segregating population derived from each of the ten wild relatives under study (**Table S5**). Each wild relative species had a subset of chromosome-specific markers validated to be polymorphic with wheat (**Table 2**). Genotyping with these chromosome-specific markers allowed differentiation of homozygous lines from heterozygous lines in the segregating populations and rapid identification of the wheat chromosome that had recombined with the wild relative segment.

The chromosome-specific markers, from homoeologous chromosomes in wheat can, collectively, detect the presence of an orthologous wild relative chromosome segment. The genotyping data shows that the markers on each of the three subgenomes for a homoeologous group, give a heterozygous call when a single wild relative segment from an orthologous group is present. However, if the orthologous segment is homozygous in the introgression lines, then the chromosome-specific markers on the wheat chromosome involved in the recombination event give a homozygous call (due to the absence of both copies of the wheat allele) while the markers on the other two subgenomes give a heterozygous call.

**Figures 4A-F** show the characterisation of introgression lines represented by the genotyping data from chromosome-specific markers alongside multicolour GISH (mcGISH) analysis of the root metaphase spreads of these lines. The distribution of chromosome-specific KASP markers, used for genotyping a wild relative species, along the 21 chromosomes of wheat is indicated by coloured regions in the bar diagrams whereas chromosomal regions lacking the presence of chromosome-specific markers for that species are indicated by white spaces. **Figures 4A-B** show the genotyping of two sister lines containing a segment(s) of chromosome 4JS of *Th. bessarabicum*. The introgression line in **Figure 4A** is heterozygous for chromosome 4JS as indicated by the presence of heterozygous calls (red regions) for diagnostic chromosome-specific markers on chromosomes 4AS and 4DS in the genotyping data (the blue regions on all the chromosomes represent markers genotyped as wheat alleles only) and validated by the presence of a single wheat-*Th. bessarabicum* recombinant chromosome in the mcGISH analysis. Chromosome 4B did not have any chromosome-specific markers polymorphic with *Th. bessarabicum* in the distal end of the short arm as indicated by a white space. **Figure 4B**, on the other hand, shows homozygous calls (green region) on chromosome 4D (alongside the heterozygous calls on chromosome 4A) indicating that the 4JS segment had recombined with chromosome 4DS of wheat and was homozygous in the line. This was validated by mcGISH which showed the presence of a homozygous chromosome T4JS-4DS.4DL. **Figure 4C** shows the genotyping of another wheat-*Th. bessarabicum* line where the markers indicate the presence of a homozygous group 5 segment from *Th. bessarabicum*, i.e. chromosome 5J, due to the presence of homozygous calls (in green) on chromosome 5A (alongside heterozygous calls (in red) on chromosomes 5B and 5D). The mcGISH analysis confirmed the presence of homozygous segment T5AS.5JL.

**Figure 4.**
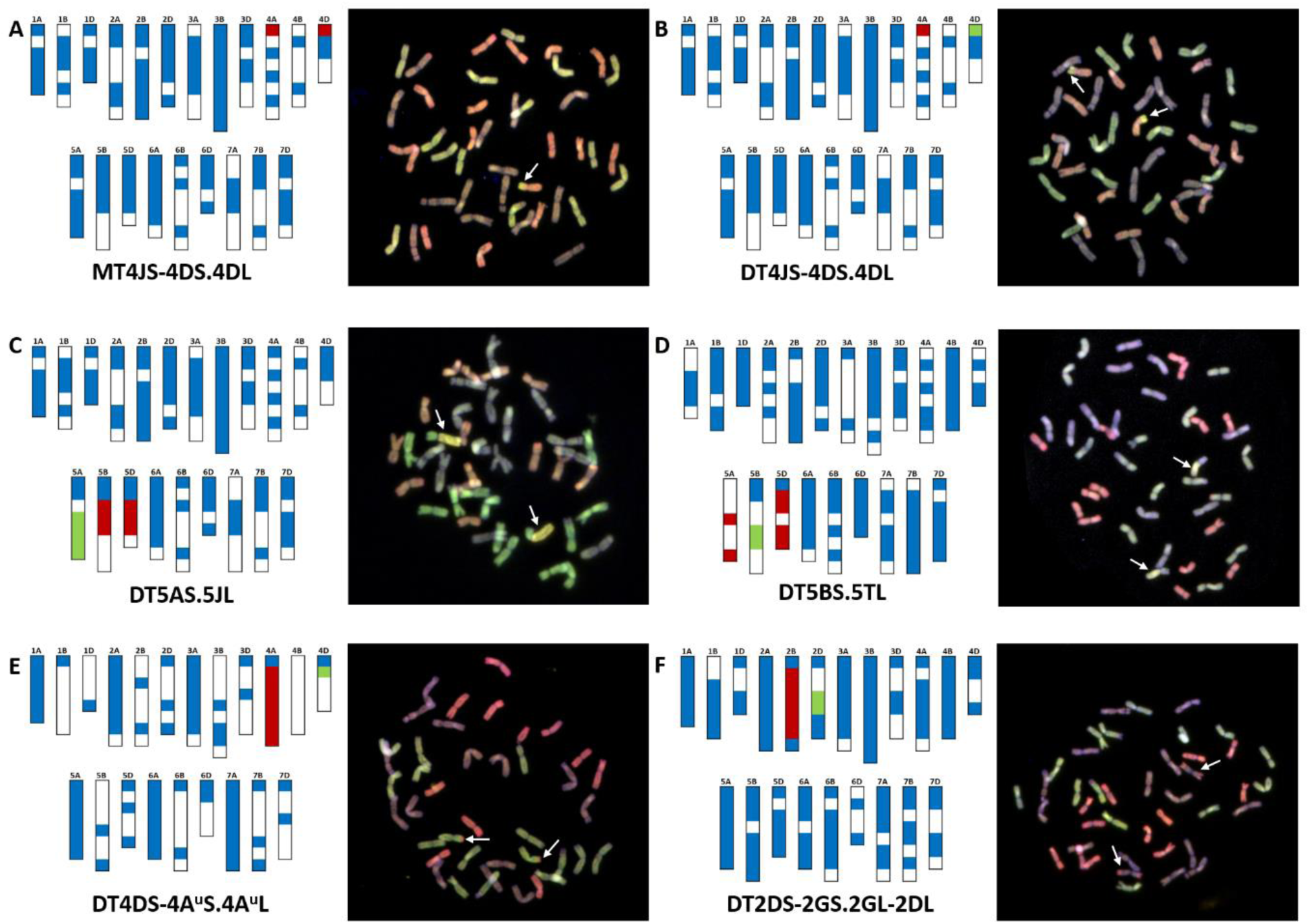
Molecular characterisation of wheat-wild relative introgression lines using chromosome-specific KASP assays and multi-colour Genomic *in situ* hybridisation (mcGISH). Genotyping data (left), using chromosome-specific KASP assays, and mcGISH analysis (right) of wheat lines carrying A) a heterozygous segment from chromosome 4JS of *Th. bessarabicum*, B) a homozygous segment from chromosome 4JS of *Th. bessarabicum*, C) a homozygous segment from chromosome 5JL of *Th. bessarabicum*, D) a homozygous segment from chromosome 5TL of *Am. muticum*, E) a large homozygous segment from chromosome 4A^u^ of *T. urartu* and F) a homozygous segment from chromosome 2G of *T. timopheevii*. In the genotyping data, all heterozygous calls are shown in red, homozygous wild relative calls in green and homozygous wheat calls in blue. White spaces indicate where there are no chromosome-specific KASP assays polymorphic between wheat and the wild relative species. Each wild relative has a species-specific set of chromosome-specific KASP assays. For the mcGISH, genomic DNA of *T. urartu* (A genome; green), *Ae. speltoides* (B genome; bluish purple) and *Ae. tauschii* (D genome; red) along with either *Th. bessarabicum* (J genome; yellow) or *Am. muticum* (T genome; yellow) were used as probes. Wild relative segments are indicated by white arrows in the mcGISH images.

The rapid detection of homozygosity and site of introgression using these chromosome-specific markers was also obtained in other wheat-wild relative introgression lines as shown in **Figures 4D-F**. The chromosome-specific markers were able to detect the homozygous presence of a wheat-*Am. muticum* recombinant chromosome T5BS.5TL which was subsequently validated by mcGISH (**Figure 4D**). The markers also detected *T. urartu* in a wheat background. **Figure 4E** shows the presence of a large *T. urartu* segment from chromosome 4A^u^ (red heterozygous calls on 4A) with markers indicating the homozygous segment had recombined with chromosome 4D of wheat (green region). Assuming most *T. urartu* chromosome segments recombine with the A genome of wheat (since *T. urartu* is the progenitor of the A genome of wheat), mcGISH is usually unsuitable for characterising wheat-*T. urartu* lines as the genomic probe used to detect *T. urartu* segments cannot differentiate between the *T. urartu* genome and the A genome of wheat. However, since the markers indicated that the *T. urartu* segment had introgressed into the D genome of wheat, mcGISH analysis could validate the presence of this segment as homozygous (**Figure 4E**). The presence of *T. timopheevii* in wheat was also easily detected by the markers as shown in **Figure 4F** where a homozygous interstitial segment from *T. timopheevii* chromosome 2G is shown to have recombined with chromosome 2D of wheat. With *T. timopheevii* being a tetraploid (2n= 4×= 28; A^t^A^t^GG), the KASP markers are not only chromosome-specific in wheat but also for the subgenomes of *T. timopheevii*. Thus, chromosome-specific markers for the A and B genomes of wheat detect the presence of the A^t^ and G genomes of *T. timopheevii* respectively. Due to the fact that there is no equivalent of the D genome in *T. timopheevii*, the chromosome-specific markers on the D genome could be polymorphic randomly with either the A^t^ or the G genomes of *T. timopheevii*. In the case of the introgression line shown in **Figure 4F**, the markers are detecting the presence of a segment of 2G via heterozygous calls on chromosome 2B (in red) and its presence as a homozygous introgression, in chromosome 2D, due to homozygous calls on chromosome 2D (in green). As with *T. urartu*, mcGISH does not usually work as a detection tool for introgressions from *T. timopheevii* in wheat. However, the detection of *T. timopheevii* is possible via mcGISH, when the markers indicate that it’s A^t^ genome has recombined with either the B or the D genome of wheat and/or it’s G genome has recombined with either the A or the D genome of wheat as shown in **Figure 4F**.

## Discussion

The Axiom Wheat Relative Genotyping Array has been used to genotype various wheat-wild relative introgression populations (Cseh et al., 2019; Grewal et al., 2018a; Grewal et al., 2018b; King et al., 2018; King et al., 2017). To cost-effectively genotype the self-fertilised progenies of these introgression lines, whilst maintaining high-throughput scale and flexibility, it was necessary to change the format of genotyping from the Axiom array to KASP assays. However, initial work to directly convert target SNP sequences into KASP assays was unsuccessful since most of the assays were detecting polymorphic homoeologous loci in wheat (data not shown). This could be likely due to the presence of homoeologous sequences that have diverged in sequence enough for probe annealing to be specific to the target subgenome during array-type genotyping while allowing amplification of homoeologous sequences by the KASP primers (possibly designed on conserved regions of the sequence). To avoid this, we focused on SNP-containing probes on the array that potentially had a single-copy in the wheat genome.

### SNP Discovery

At the time of the initial BLASTN search, the assembly used for the analysis was the IWGSC Chinese Spring CSS v3 (IWGSC, 2014). Approximately 7.4% of the probes on the array were identified as potentially being in single-copy regions in wheat. Primers were designed to amplify these regions to obtain more information on the extended flanking sequences of the SNPs. Approximately 80% of the identified probes were able to have primers designed from their flanking sequence since not all primer sequence combinations complied with optimum design parameters. In order to find SNPs between wheat and ten wheat relatives (currently under study at the Nottingham BBSRC Wheat Research Centre), it was necessary that the primers amplified at least one of the two wheat varieties used for PCR, along with one or more of the ten wild relative species. From the 2,170 primer pairs designed, 79% were successful at such amplification and the resulting PCR products were sent for sequencing. A second PCR amplification attempt was made for every failed primer, but it is possible that due to sub-optimal conditions some chromosomes were more successful at amplification than others. Multiple sequence alignment of wheat and its wild relatives yielded 2,374 putative SNPs from 8,451 sequences spread across the 21 chromosomes of wheat (**Table 1 and Table S1**).

### Primer design for subgenome-specific assays

To verify that SNPs obtained after PCRs and sequence analysis were in single-copy regions of the wheat genome, a second BLASTN search was conducted using the sequenced amplicons against the new IWGSC wheat genome sequence Refseq v1 (IWGSC et al., 2018). The results showed that less than one-fifth of the sequences belonged to single-copy regions, with most having 3 homoeologous copies in wheat (**Table S2**). This is potentially due to the difference between the quality of the two genome assemblies used for the BLASTN searches, since the key distinguishing feature of the IWGSC RefSeq v1 is that it is an assembly of long-reads, with 90% of the genome represented in superscaffolds larger than 4.1 Mb (IWGSC et al., 2018), making it a more reliable, high quality reference assembly for wheat.

It was possible to annotate sequences to allow ‘anchoring’ of the common primer to a subgenome-specific base, thereby optimising the primer design to produce target-specific KASP assays as shown in **Figure 1B**. Previous studies have used this technique successfully to design chromosome-specific KASP assays in wheat (Allen et al., 2011) but also indicated that in the absence of a software that would automatically annotate sequences with anchored bases, it was a time-consuming process. However, the lack of availability of both the wild relative genome sequences and a complete wheat genome sequence made the approach taken in this study the most appropriate at the time. In future, automated pipelines such as PolyMarker™ (Ramirez et al., 2015) and MAGICBOX™ (Curry et al., 2016) will be very useful tools to redesign failed assays or design chromosome-specific assays for newly discovered SNPs between wheat and its wild relatives.

### SNP validation and characterisation

A subset of 1,000 putative interspecific SNPs was selected for conversion to KASP assays (**Table 1 and Table S3**). After genotyping wheat varieties, wild relative accessions, Chinese Spring nullisomic lines and various back-cross populations (**Table S4 and S5**), the results showed that 73% of the SNPs were converted into a working KASP assay. This conversion rate is lower compared to another study in which 96% were successfully validated to be polymorphic between wheat varieties as described by Allen et al. (2013) but still relatively high for a complex polyploid such as wheat (Edwards et al., 2009). However, it was noted that approximately half the failed assays amplified the wheat varieties but not the target wild relative accessions (**Data S1**). This could be possibly due to several reasons such as sequencing errors leading to false positives during SNP discovery, inefficient primer design for the wild relative allele and/or suboptimal PCR conditions during genotyping. Of the assays that worked, 17 were found to be polymorphic within wheat probably due to the presence of homoeologous sequences that were not detected in the sequence data but were amplified by the KASP primers.

Most of the working assays for a target chromosome were able to distinguish the heterozygous samples from the homozygous samples in a segregating population and provided a null call in the corresponding Chinese Spring nullisomic tetrasomic line, and were thus classified as chromosome-specific (**Figures 2A-E**). From amongst the validated assays, 620 were chromosome-specific (**Table 2**) and 90 were chromosome-nonspecific i.e., they detected more than one homoeologous loci in wheat (**Figures 2F-H**). Various wild relative species had validated chromosome-specific assays that also worked in other species (**Table S6**), with more than half shown to be working for at least 3 wild relative species (**Data S2**), thereby demonstrating the diverse applicability of these assays for various wheat-wild relative breeding programmes. BLASTN results showed that ∼62% of these chromosome-specific assays were derived from single-copy regions of the wheat genome (**Data S3**). It is possible that such regions were either unique to only one progenitor genome or one or more copies could have been lost after polyploidisation. BLASTX results showed that 275 (44%) of the chromosome-specific assays were in protein coding regions with ∼52% of these being single-copy loci in wheat (**Data S3**). Previous studies have hypothesised that codominant SNP assays are most likely to be in single-copy genes of as yet unknown function in wheat. Where they are found to be in 3-copy genes it is likely to be in 3’ UTR regions which are more divergent than protein-coding sequence (Allen et al., 2013). In our study, these single-copy regions were found in both landraces such as Chinese Spring, and modern cultivars such as Paragon, Pavon 76 and Highbury. Thus, it is possible that these contigs represent genes that were lost before or during the domestication process. Previous reports have documented intra-or intervarietal heterogeneity and gene loss within elite or inbred lines of wheat (Tokatlidis et al., 2004; Winfield et al., 2012).

BLASTN results of the chromosome-specific (**Data S3**) and the nonspecific (**Data S4**) SNP sequences allowed the visualisation of the distribution of all the KASP markers (**Figure 3**), diagnostic for the ten wild relatives used in this work, and identification of regions where gaps exist and need to be filled in the future through more SNP discovery, KASP assay design and validation.

### Genotyping with chromosome-specific markers

Through the Axiom^®^ Wheat-Relative Genotyping array, it was possible to detect the presence of wild relative segments in a wheat background in various back-crossed populations. However, when the lines needed to be self-fertilised to create stable introgression lines, the array was unable to effectively distinguish between heterozygotes and homozygotes. Moreover, genotyping with the array could not provide any information about which specific wheat chromosome had recombined with the wild relative species. The development of the chromosome-specific KASP markers has, for the first time, allowed the identification of homozygous introgressions and the site of recombination in wheat.

In cases where the introgressed segment from a wild relative chromosome is likely to be orthologous with the wheat chromosomes, its presence is indicated by heterozygous calls for chromosome-specific markers on homoeologous loci across all three subgenomes of wheat (**Figure 4A**). This is because the markers were designed to be polymorphic between the wild relative genome and each of the three subgenomes within a homoeologous group. However, when the recombinant segment is homozygous in wheat, the loss of wheat alleles (due to both copies of the wheat loci on one subgenome being replaced by wild relative loci) results in a homozygous wild relative call for the chromosome-specific markers on the recombinant wheat chromosome (**Figure 4B**) and hence allows for the identification of the site of introgression.

False positives of homozygosity could be obtained if there is a deletion of both copies of a wheat subgenome from a homoeologous group which is the same as the one into which the wild relative segment has been introgressed. Gaps in the marker distribution, particularly in the distal regions of chromosomes, might result in difficulty in distinguishing between a large recombinant segment and a whole chromosome introgression and also in the failure to detect small telomeric introgressions. Another point to note is that these markers were designed assuming overall macro-synteny between the wheat subgenomes and the wild relative genomes. However, there are wild relative genomes with major rearrangements compared to the wheat genome, such as *S. cereale* (Devos et al., 1993; Li et al., 2013), in which case known rearrangements must be taken into account. For less well characterised wild relatives, it will still be possible to use the markers to identify the presence of wild relative segments and to distinguish between heterozygous and homozygous introgressions.

## Conclusion

This study has described the design, validation and implementation of chromosome-specific KASP markers in wheat. A majority of these markers are based on single-copy regions in the wheat genome but where there are homoeologous copies of the target SNP sequence, ‘primer anchoring’ was used to design chromosome-specific assays. Thus, 620 chromosome-specific KASP assays have been validated which allow the rapid identification of homozygous wild relative introgressions in a wheat background and their potential site of recombination within wheat. In addition to this, 90 chromosome-nonspecific KASP markers were also identified which can be used for the detection of wild relative chromatin in introgression lines. Most of the developed assays can be used for detection of multiple wild relative species used in this study. Thus, there is potential for these markers to be used to detect the presence of various other wild relative species and moreover, for the detection of wild relative introgressions in a durum background. As such, these KASP assays could be a highly valuable resource, which will be of considerable interest to wheat researchers and, in particular, the breeding community.

## Experimental Procedures

### Plant Material

The wheat varieties and accessions of ten wild relatives (**Table S4**) were grown for leaf tissue collection and nucleic acid extraction. The whole set of Chinese Spring nullisomic tetrasomic lines were obtained through the Germplasm Resource Unit (John Innes Centre; www.seedstor.ac.uk). The back-cross populations, created from crossing each of the wild relatives with the wheat cv. Paragon, were generated at the Nottingham BBSRC Wheat Research Centre.

All plants were grown in pots in John Innes No. 2 soil and maintained in a glasshouse at 18–25 °C under 16 h light and 8 h dark conditions. Leaf tissues were harvested from 3-week-old plants. All harvested tissues were immediately frozen on liquid nitrogen and stored at −80 °C until nucleic acid extraction.

### Nucleic Acid Extraction

Genomic DNA was extracted according to the Somers and Chao protocol (http://maswheat.ucdavis.edu/PDF/DNA0003.pdf, verified 21 January 2019, original reference in Pallotta et al., 2003). In case of wild relatives with multiple accessions, the genomic DNA was pooled into one sample.

### Primer Design

All SNP probe sequences on the Axiom^®^ Wheat-Relative Genotyping Array were used in a BLASTN search (e-value cut-off of 1e-05) against the wheat reference sequence (IWGSC CSS v3; IWGSC, 2014) to find probes that had a BLAST hit to only one contig in the wheat genome. Primers were designed from the flanking 500 bp sequence using Primer 3 v4.1.0 (Untergasser et al., 2012) with default primer size and Tm conditions. Primers were ordered through Eurofins Genomics, Germany.

### Polymerase Chain Reaction (PCR) and Sequencing

All primers were used for PCR amplification of genomic DNA using a touchdown program on the Mastercycler nexus GSX1 (Eppendorf, Germany): 95°C for 5 min, then 10 cycles of 95°C for 1 min, 65°C for 30 s [–1°C per cycle] and 72°C for 2 min, followed by 40 cycles of 95°C for 1 min, 58°C for 30 s, and 72°C for 2 min. The amplification products were run on a 1.5% agarose gel with size marker Hyperladder™ 1kb (Bioline, UK). DNA bands (∼ 500 bp) were cut from the gel, cleaned using the NucleoSpin Gel and PCR Clean-up kit (Macherey-Nagel, Düren, Germany) and sent for Sanger sequencing (Source Biosciences, Nottingham, UK).

### SNP Discovery

Sequences were visualised using Chromas Lite v2.1.1 (Technelysium, Australia). All sequences from the same primer pair were aligned using GeneDoc v2.7. The Chinese Spring sequence, for each primer pair, were used in a BLASTN search (e-value cut-off of 1e-05) against the new wheat reference sequence (IWGSC RefSeq v1; IWGSC et al., 2018) to check for homoeologous sequences and obtain the physical position of the target SNP. The target interspecific SNP was annotated by its IUPAC code and square brackets. Any other SNPs found in the 100 bp region flanking the target SNP were also annotated with the corresponding IUPAC code. If a SNP-containing sequence had more than one homoeologous copy in wheat, then any subgenome-specific bases, for the target subgenome, in the 100 bp sequence flanking the SNP were annotated with chevrons.

### KASP™ Assay Design and Validation

For each putative SNP, KASP™ assays containing two allele-specific forward primers and one common reverse primer (**Data S5**) were designed (LGC Genomics, Middlesex, UK) using the annotated SNP sequences. Leaf tissues from all the ten backcross populations (**Table S5**), the parental lines and the Chinese Spring nullisomic lines were sent for DNA extraction and genotyping with the KASP™ assays (LGC Genomics, Middlesex, UK).

### Multi-colour Genomic *in situ* Hybridisation (mcGISH)

Preparation of the root metaphase chromosome spreads, the protocol for the mcGISH and the image capture was as described in King et al. (2017). All slides were probed with labelled genomic DNA of the three putative diploid progenitors of bread wheat, i.e. *T. urartu* (A genome), *Ae. speltoides* (B genome), and *Ae. tauschii* (D genome). Additionally, introgression lines with segments from *Th. bessarabicum* and *Am. muticum* were probed with the respective wild relative’s labelled genomic DNA. The genomic DNA of 1) *T. urartu* was labelled by nick translation with ChromaTide™ Alexa Fluor™ 488-5-dUTP (Invitrogen; C11397; coloured green), 2) *Ae. speltoides* was labelled by nick translation with DEAC-dUTP (Jena Bioscience; NU-803-DEAC; coloured blueish purple), 3) *Ae. tauschii* was labelled with ChromaTide™ Alexa Fluor™ 594-5-dUTP (Invitrogen; C11400; coloured red) and 4) *Th. bessarabicum* and *Am. muticum* were labelled by nick translation with ChromaTide™ Alexa Fluor™ 546-14-dUTP (Invitrogen; C11401; coloured yellow).

## Acknowledgements

This work was supported by the Biotechnology and Biological Sciences Research Council (Grant No. BB/P016855/1) as part of the Developing Future Wheat (DFW) programme. The funding body played no role in the design of the study and collection, analysis, and interpretation of data and in writing the manuscript.

## Supporting Tables

Table S1 Distribution of the number of SNPs discovered for each wild relative species, on the wheat chromosomes.

Table S2 Distribution of second BLASTN results, for the discovered SNPs, across each wheat chromosome.

Table S3 Distribution of the number of SNPs selected for KASP assay design, for each wild relative species, on the wheat chromosomes.

Table S4 Details of the parental lines used for SNP discovery.

Table S5 Number of lines in each backcross population, for a wild relative species, used for genotyping.

Table S6 Number of chromosome-specific KASP assays common between two wild relative species.

## Supporting Data

Data S1 Genotypes of all the parental and control lines with working KASP assays.

Data S2 Details of wild relative species validated for each polymorphic KASP assay.

Data S3 BLASTN and BLASTX results for all chromosome-specific SNPs.

Data S4 BLASTN results for all chromosome-nonspecific SNPs.

Data S5 KASP assay primer details for all selected SNPs.

